# Inference and analysis of population structure using genetic data and network theory

**DOI:** 10.1101/024042

**Authors:** Gili Greenbaum, Alan R. Templeton, Shirli Bar-David

**Affiliations:** Department of Solar Energy and Environmental Physics, Blaustein Institutes for Desert Research, Ben-Gurion University of the Negev, Sede Boqer Campus 84990, Israel; Mitrani Department of Desert Ecology, Blaustein Institutes for Desert Research, Ben-Gurion University of the Negev, Sede Boqer Campus 84990, Israel; Department of Biology, Washington University, St. Louis, MO 63130, USA; Department of Evolutionary and Environmental Ecology, University of Haifa, Haifa 31905, Israel

## Abstract

Clustering individuals to subpopulations based on genetic data has become commonplace in many genetic studies. Inference of population structure is most often done by applying model-based approaches, aided by visualization using distance-based approaches such as multidimensional scaling. While existing distance-based approaches suffer from lack of statistical rigor, model-based approaches entail assumptions of prior conditions such as that the subpopulations are at Hardy-Weinberg equilibria. Here we present a distance-based approach for inference of population structure using genetic data by defining population structure using network theory terminology and methods. A network is constructed from a pairwise genetic-similarity matrix of all sampled individuals. The *community partition*, a partition of a network to dense subgraphs, is equated with population structure, a partition of the population to genetically related groups. Community detection algorithms are used to partition the network into communities, interpreted as a partition of the population to subpopulations. The statistical significance of the structure can be estimated by using permutation tests to evaluate the significance of the partition’s *modularity*, a network theory measure indicating the quality of community partitions. In order to further characterize population structure, a new measure of the Strength of Association (SA) for an individual to its assigned community is presented. The Strength of Association Distribution (SAD) of the communities is analyzed to provide additional population structure characteristics, such as the relative amount of gene flow experienced by the different subpopulations and identification of hybrid individuals. Human genetic data and simulations are used to demonstrate the applicability of the analyses. The approach presented here provides a novel, computationally efficient, model-free method for inference of population structure which does not entail assumption of prior conditions. The method is implemented in the software NetStruct, available at https://github.com/GiliG/NetStruct.

## 1 Introduction

Inference and analysis of population structure from genetic data is often used to understand underlying evolutionary and demographic processes experienced by populations, and is an important aspect in many genetic studies. Such inference is mainly done by clustering individuals into groups, often referred to as demes or subpopulations. Evaluation of population structure and gene flow levels between subpopulations allows inference of migration patterns and their genetic consequences [1, 2]. As sequencing of larger portions of the genome is becoming more readily available, there is an increasing need for a variety of computationally-efficient statistically-testable methods for such inference.

Analysis of population structure can be done at the subpopulation-population level by assuming putative subpopulations and studying how these relate genetically (e.g. F-statistics, AMOVA[3], phylogenetic methods [4, 5, 6]), or at the individual-subpopulation level by attempting to cluster individuals to subpopulations. The methods for clustering individuals based on genetic data can be further divided to two categories: Model-based approaches and distance-based approaches[7, 8, 9]. Model-based approaches evaluate the likelihood of the observed data assuming that they are randomly drawn from a predefined model of the population, for example that there are *K* subpopulations and that these subpopulations are at Hardy-Weinberg equilibrium (HWE). Distance based approaches aim at identification of clusters by analysis of matrices describing genetic distances or genetic similarities between individuals or populations, for example by visualization using multidimensional scaling (MDS) methods such as Principle Component Analysis (PCA). Distance-based methods are ususally model-free and do not require prior assumptions, as with the model-based methods. Over the last decade or so, model-based methods have been more dominant as procedures for inference of population structure, mostly with implementation of Bayesian clustering and maximum-likelihood techniques in programs such as `STRUCTURE`[7], `ADMIXTURE`[8] and `BAPS`[10]. It has been pointed out that distance-based methods have several disadvantages: they are not rigorous enough and rely on graphical visualization, they depend on the distance measure used, it is difficult to assess significance of the resulting clustering, and it is difficult to incorporate additional information such as geographical location of the samples[7] ([11] and [12] address this last concern). Given these disadvantages, it would seem that distance-based measures are less suitable for statistical inference of population structure. On the other hand, model-based approaches suffer from the necessity to restrict the interpretation of the results by heavily relying on the prior assumptions of the model, for example that the populations meet certain equilibria conditions, such as migration-drift or HWE[7].

There has recently been a flourish of network theory applications to genetic questions in genomics[13], landscape genetics[14], migration-selection dynamics[15], and population structure at the subpopulation-population level [16, 17, 18, 19]. Recently, a network-based visualization tool (`NETVIEW`[20]) of fine-scale genetic populations structure, using a Super Paramagnetic Clustering algorithm[21], has been proposed and successfully applied to analysis of livestock breeds[22, 23], and other network clustering approaches have also been implemented on genetic data[24]. However, these methods still suffer from the many disadvantages of distance-based clustering approaches, and a more rigorous and statistically testable distance-based approach is still missing.

Development of a suitable distance-based network approach, that will coherently address inference of population structure from genetic data and not suffer from the disadvantages listed above, necessitates a clear definition of genetic population structure in equivalent network theory terminology. A genetically defined subpopulation is commonly thought of as a group of individuals within the population which are more genetically related (or more genetically similar) to each other than they are to individuals outside the subpopulation, as a result of many possible genetic processes such as genetic drift, migration, mutation and selection. In a network, a group of nodes which are more densely and strongly connected within the group than outside the group, relative to the given topology of the network, is called a *community*[25]. Therefore, in network theory terminology, the equivalent of a genetic population structure should be the community partition of a network constructed with individuals as nodes and edges defined using an appropriate genetic distance or similarity measure. In network science, clustering nodes into groups has been extensively studied, and specifically community detection has attracted much interest[26]. Since there is no single rigid definition of a community, and identifying optimal partitions is computationally expensive, many approaches and algorithms to optimally detect communities in networks have been proposed[27,28].

We propose a network-based approach for analyzing population structure based on genetic data. We show that by applying recent advances in network theory, it is possible to design a distance-based approach that overcomes the limitations of other distance-based approaches, and does not suffer from the disadvantages of model-based approaches. We also show how rigorous statistical inference can be incorporated into this network-based approach in a manner that does not entail prior assumptions or conditions about the data.

The process can be used with a large number of loci (e.g. microsattelites, SNPs) since it is computationally efficient in regards to the number of loci incorporated in the analysis. Moreover, we define a new measure for the strength to which an individual is associated with its assigned community, called Strength of Association *(SA)*, and we show how Strength of Association Distribution (SAD) analysis can be used to infer further details regarding population structure, such as gene flow patterns of each subpopulation and identification of potentially hybrid individuals. The analysis is demonstrated on genetic data from human population extracted from the HapMap project[29], as well as on simulated data. In addition to presentation of a new distance-based method for population structure inference, we believe that defining the problem of genetic population structure analysis in network terminology will allow future adoption and adaptation of network methods and techniques to address population genetic questions.

## 2 Methods

In this section we provide the relevant theory and describe a network-based approach for constructing genetic networks and inferring population structure by detecting community partitions in these networks. Following detection of community structure, we propose an additional exploratory analysis, based on a measure of the strength of association of individuals to communities, that may shed light on finer details of the community structure and therefore on the population structure.

### 2.1 Constructing networks from genetic data

A network is a set of discrete entities, *nodes*, where each pair of nodes may be connected by an *edge*, possibly characterized by a *weight*. Networks can be described by adjacency matrices, where the element in column *i* and row *j* is the weight of the edge connecting node *i* and node *j*. For most network applications, dyads connected by edges with high-valued weights are interpreted as being strongly connected. Therefore, a genetic-similarity matrix (a matrix describing some measure of genetic similarity or relatedness between all pairs of individuals, based on their genotypes) of a population can be regarded also as the adjacency matrix of a genetic-similarity network. The more classic genetic-distance (genetic dissimilarity) matrices are similar, only that in these high values indicate weakly connected dyads, and therefore such matrices must be appropriately transformed in order to be considered as network adjacency matrices. There is no fundamental difference, for our purpose, between similarity and distance matrices. Many genetic similarity, distance and relatedness measures have been proposed[30], but if we restrict the discussion to symmetric similarity measures, where similarity between individual *i* and *j* is the same as between individual *j* and *i*, the genetic network thus described is a weighted undirected network, where each edge is characterized by a weight but does not have directionality. Since we would like to consider not only allele sharing between individuals, but also differences in allele frequencies between subpopulations, we further restrict the discussion to genetic similarity measures which are expressed relative to allele frequencies in a reference population, i.e. measures that incorporate the allele frequencies of the total sampled population. These measures should not incorporate allele frequencies other than those of the total sample (e.g. allele frequencies in sample sites or other locally-defined sampled populations), since this would mean that the null hypothesis is other than that there is no population structure. From a distance-matrix perspective, the null hypothesis is that the adjacency matrix, appropriately transformed, is a symmetric individual-pairwise distance matrix, such that all individuals are equally distant from the multi-loci centroid of the total population, with no sub-clustering.

In a network such as described above, the strength of the connection between each dyad is relative to the genetic similarity between them, where shared rare alleles convey a stronger connection than do common alleles. This is different from many commonly-used distance measures for construction of individual-level distance matrices, such as in AMOVA [3], where distances are measured by mismatch of alleles, irregardless of their frequencies. Since even unrelated individuals may share many alleles, especially when many loci are examined, it is likely that this network will be extremely dense. It may therefore be useful, both from a computational point-of-view and in order to emphasize strong genetic relations within the population so as to increase detection power of network procedures, to remove edges which describe weak connections. This can be done in different ways, but the most straightforward approach is to remove edges with weights below a certain threshold, which is the approach we implement here. In this way a sparser network that consists of strong relatedness interconnections is attained.

Since using different thresholds will result in different networks which may give, for the analyses described below, different population structures, we recommend systematically exploring different threshold values. For very low threshold values many weak relatedness interconnections will be included in the network, which may result in very dense networks which could mask related groups within the population. Very high thresholds may result in the network breaking down into many disconnected components (a network component is a group of nodes that are connected within themselves but are not connected to any other node in the network), up to a point when the network includes only very small groups of connected nodes. Such networks are most likely not informative of population structure since they represented too few related dyads, and the community partition will likely consist of many one- or two-node communities. Each community is confined to be within a component, and if the network consists of many small components then the community partition is constrained to include many small communities. Therefore the informative structures should be detected at the intermediate thresholds, and different thresholds in this range may describe structure at different hierarchical levels *(see Analysis of human SNP data* section for an example of a systematic exploration of threshold values).

### 2.2 Network communities and genetic population structure

In network theory, the term *community* refers to a subset of nodes in a network more densely connected to each other than to nodes outside the subset [31]. There are now several algorithms for efficiently partitioning a network into communities [27, 28]. Most commonly, a partition of a network into communities is evaluated by calculating the *modularity* of the partition, a quality measure (between -1 and 1) indicating whether the partition is more or less modular than would be expected if connections were randomly distributed[32]. The modularity of a particular community partition of a weighted network A (a network with weights assigned to its edges) is defined as the weight of the intra-community connections minus the expected weight of the intra-community connections in a random network preserving the edge weights of each node[33]:

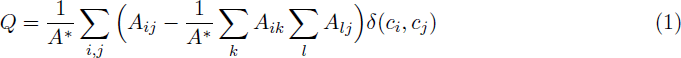

where 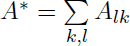 is the sum over all edge weights in the network and 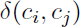 is a delta function with value one if nodes *i* and *j* are in the same community and zero otherwise. A positive modularity value indicates that the partition is more modular than expected. Many community detection algorithms try to approximate an optimal partition, in which the modularity measure is maximal over all possible partitions. A partition of one community, which includes all nodes in the network, results in a modularity of zero, and therefore for every network the optimal partition, maximizing the modularity, is always non-negative. Community detection algorithms do not generally require a priori knowledge of the number of clusters present in the network.

Since in a genetically subdivided population the individuals in a subpopulation are expected to be more highly related in comparison to a random redistribution of relatedness levels between individuals, communities in the genetic network are expected to coincide with the subpopulations of the underlying population structure. We therefore propose that population structure can be ascertained by constructing a genetic network based on a genetic similarity measure, and then applying community detection methods to identify a partition which maximizes modularity. Since it is possible that an optimal partition consists of just one community (i.e. the entire network), community detection algorithms can also identify scenarios with no population subdivision.

A partition of the network into two or more communities may be indicative of population structure, but community partitions may also be detected in panmictic populations due to chance alone. In order to test whether the detected population structure is significant, we test whether it is significantly different than the null hypothesis of no population structure. Several approaches have been suggested in order to evaluate the statistical significance of community partitions [27], but here we pursue a significance test based on permutations of the genetic network, which focuses on testing whether the null hypothesis can be rejected or not. With no population structure, dense subgraphs are not expected to appear in networks constructed as described above, and if such subgraphs do appear, the modularity of an optimal partition of the network should be relatively low. In randomly permuted networks, dense subgraphs are expected to be present as often as in unstructured populations, and hence optimal partitions of permuted networks can be used to represent the expected modularity of optimal partitions of unstructured populations. If the modularity of the detected community partition is not significantly higher than modularities of optimal partitions of the permuted networks, we cannot reject the null hypothesis. We can, therefore, use permutations to test the statistical significance of detected community partitions.

With the application of this significance test, population genotypes are considered to describe a structured population if, and only if, more than one community is detected, and the detected partition is more modular than would be expected in a population with no structuring. Note that the permutation can be done at the level of the genotypes, or, if the network is large enough, at the adjacency matrix level, with symmetry-preserving permutations of the matrix. Permutations of the genotypes require constructing many matrices, which would be computationally expensive when many loci are involved. Therefore permuting the adjacency matrix should be considered as an alternative when the network contains many edges, since in large networks weight values should not constrain the possible attainable modularity values of optimal partitions.

### 2.3 Strength of Association Distribution (SAD) analysis

Revealing the division of the population into subpopulations may shed light on many aspects of the underlying evolutionary and ecological processes; however, more information can be attained by further analyzing the characteristics of the partition. The partitioning of the network into dense subgraphs, as presented above, does not convey information regarding how important each individual is to the detected partition. Here we introduce a measure intended to evaluate this aspect, the *Strength of Association (SA)* of individual *i* to its community. Given a community partition *C* and an individual *i*, we define the Strength of Association as

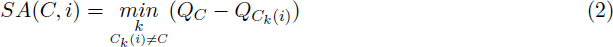

where *Q_C_* is the modularity of the partition *C* and *C_k_*(*i*) is the partition identical to *C* except that node *i* is assigned to community *k* instead of its original community. Thus, high *SA* values indicate that the partition *C* is sensitive to the assignment of *i*, and that the assignment of *i* to its community is essential, whereas low *SA* values indicate that there is another community that the individual is well assigned to. From a population genetic perspective, the measure evaluates how strongly individuals are related to the group to which they were assigned to, and *SA* is expected to be low when individuals are recent descendants from individuals from more than one subpopulation. For example, potential hybrids are expected to show low *SA* values, and the *k* that minimizes equation 2 is the probable origin of the second lineage of the individual (see *Identifying hybrids and recently admixed individuals* section for an example).

The *SA* measure is a measure at the individual level, taking into account genetic data of the entire population. We introduce an exploratory analysis that evaluates characteristics of subpopulations relating to the cohesion of the subpopulation and the association of individuals to the subpopulation, the *Strength of Association Distribution* (SAD) analysis. This analysis examines the distribution of the *SA* values of the different communities and compares the statistical attributes of these distribution (e.g. the mean, variance and skew of the *SA* distributions). Since different scenarios are expected to result in different cohesion of the subpopulations, it may be possible to hypothesize what underlying processes were responsible for shaping the genetics of the population.

For example, a closed disconnected subpopulation is expected to display a narrow SAD with high mean (high community cohesion), since in a closed population individuals will be strongly related relative to the entire population, and individuals descended from lineages outside the subpopulation are rare. A subpopulation experiencing constant moderate gene flow levels is expected to display a wide or left-skewed SAD with high mean, since there should be many individuals with lineages that are mostly from the subpopulation, but recent migrant and descendants of recent migrants are expected to have low *SA* values, increasing the variance and the left-skewness of the distribution. A subpopulation experiencing constant strong gene flow levels is expected to display a SAD with low mean, as many individuals will be descendants of migrants.

## 3 Analysis of human SNP data

In order to demonstrate the applicability of the network approach to infer population structure and the SAD analysis, we have selected a data set of human SNP data, extracted from the Hapmap database [29]. This data set is well suited for the demonstration of a new approach since it is taken from a population where structure and demographic history are well known from archaeological, historic and genetic studies. The genetic data for this analysis consisted of 50 randomly selected individuals from each of the 11 sampled populations of the HapMap project (overall 550 individuals): African ancestry Americans (A1), Africans from west Kenya (A2), Masai Africans (A3), Africans from Nigeria (A4), Europeans from Italy (E1), European ancestry Americans (E2), Han Chinese (C1), Chinese ancestry Americans (C2), Japanese (J), Indian ancestry Americans (I) and Mexican ancestry Americans (M). For each individual, 1000 polymorphic SNPs from each autosome were randomly selected (overall 22,000 sites per individual). In order to compare the results with a modelbased approach, the same data were analyzed with the most widely-used model-based software, `STRUCTURE` [7].

### 3.1 Network construction

A genetic network was constructed from the genotypes (without any information on the original grouping of the individuals) using, for calculation of genetic similarity, a simple symmetric frequency-weighted allele-sharing similarity measure. Analogous to the molecular similarity index[30, 34], we defined the frequency-weighted similarity at locus *l* for diploid individual *i* with alleles *a* and *b*, with frequencies *f_a_* and *f_b_* (in the total sample) respectively, and individual *j* with alleles *c* and *d*:

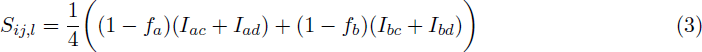

where *I_ac_* is one if alleles *a* and *c* are identical and zero otherwise, and the other indicators similarly defined. Note that this measure is a multi-allelic measure, and is commutative with respect to *i* and *j*. Given a sample with *L* loci, the weight of the edge connecting individuals *i* and *j* is defined as the mean frequency-weighted similarity over all loci:

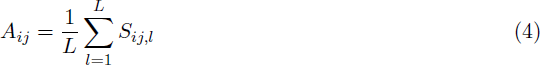

The relatedness measure defined in equation 3 is a very simple symmetric relatedness measure, that measures diversity relative to the entire population, since it takes into account the allele frequencies at the level of the entire population (with sharing of rare alleles conveying a stronger connection than sharing common ones). Other, more sophisticated, measures are likely to construct more accurate networks and may be specific to the type of marker considered (e.g. for microsatellites the length of the repeat might be taken into account) or include additional information (e.g. geographic locations of the samples). The formulation presented here is designed to analyze diploid populations, but it can be easily generalized to any level of ploidy.

### 3.2 Community partition

There are currently many algorithms used for detecting population structure, relying on different network theory concepts (reviewed by Fortunato[27] and Lancichinetti[28]). We have used several of the commonly used algorithms (implemented using igraph[35]), presented in Appendix A, and we show here the results from the classic Girvan-Newman algorithm[26] (the results using different algorithms are not qualitatively different).

The edge-removal threshold parameter was systematically explored by examining thresholds from 0 until all edges were removed, in steps of 0.001. For very low thresholds (0-0.181) the constructed networks were very dense and no population structure was detected (only one community was found which included all nodes). As mentioned above, this is to be expected from all but the most distinctly structured populations, since including connections between very weakly related individuals decreases the capability of the community detection algorithms to detect dense sub-graphs within the network. For very high thresholds (above 0.209) the networks break up into many disconnected components, many of which include only one or two nodes. Such networks cannot be coherently analyzed for communities (see *Methods* section).

For thresholds between 0.182 and 0.208, different community partitions were detected for different threshold ranges (Fig. 1). For thresholds in the range 0.182-0.188 two communities were detected, and Figure 1C shows results for threshold 0.188, referred to as “low threshold”. For the range 0.189-0.195 three communities were detected, and Figure 1B shows results for threshold 0.194, referred to as “medium threshold”. For thresholds above 0.196 the network was no longer connected and broke into several components, most notably a dense East Asian component and the rest of the network composed of one or more components. For the range 0.196-0.200 one community was detected in the East Asian component and four communities in the rest of the network. For thresholds above 0.201 only the East Asian component remained intact while the rest of the network broke into many small components and could no longer be meaningfully analyzed. The East Asian component consisted of one community for the threshold range 0.196-0.206 and two communities for the threshold range 0.207-0.208. Figure 1A shows the results of the community partition with threshold 0.207 of the East Asian component, with two communities, and for threshold 0.198 for the rest of the network, with four communities (referred to as “high threshold”). Within the edge-removal threshold ranges mentioned above, there was no significant change in the assignment of the individuals to communities. Therefore, three qualitatively different community partitions of the network into communities have been found by systemically testing different edge-removal thresholds, with either two, three or six communities for low, medium and high thresholds respectively (Fig. 1).

**Figure 1.**
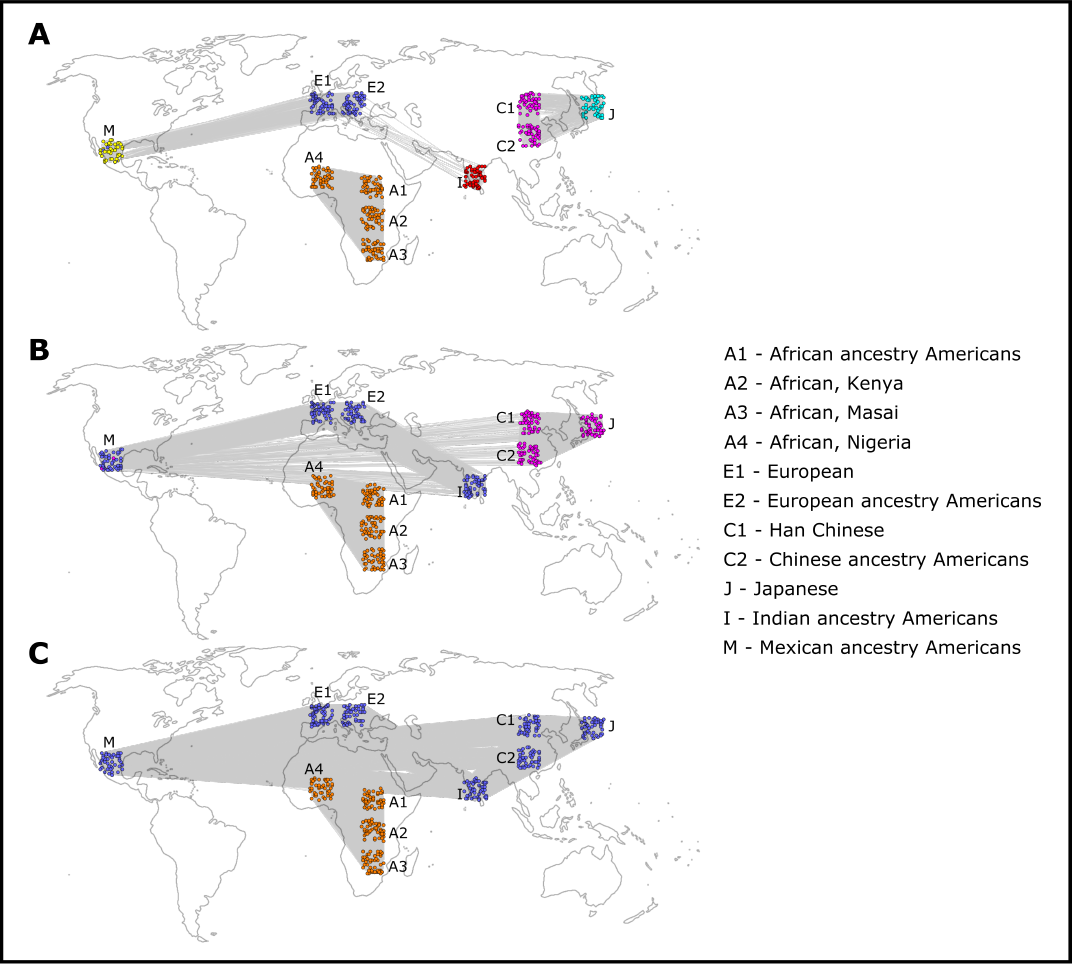
Community detection on three networks with different thresholds. Each node represents an individual, with colors representing the community assigned by the community detection algorithm. (A) high threshold (0.207 for East Asian component, 0.198 for the rest of the network) (B) medium threshold (0.194) (C) low threshold (0.188). For visualization purpose, individuals are placed on the world map roughly corresponding to their ancestry.

Permutation tests using 1000 permutations of the genotypes were conducted, and all detected community partitions were strongly significant (*p* ≤ 0.001). With the low threshold the partition corresponded to an African\Non-African division (Fig. 1C), with the medium threshold to an African\Indo-European\East Asian division (Fig. 1B), and with the high threshold to one of six communities: African, Indian, European, Mexican, Chinese and Japanese (Fig. 1A; Some of the other community-detection algorithms also detected the Masai population as a distinct community for the high threshold, Appendix A). The trend where higher thresholds reveals more detailed structure is correlated with the historically known patterns of human population differentiation. The low threshold coarse division of the population into two groups corresponds with the more ancient “out-of-Africa” migrations, the medium threshold division of the Eurasian population corresponds with the more recent migrations to Asia and the long-lasting Indo-European ethnic ties, and lastly the high thresholds correspond with the most recent events of migration and isolation leading to the formation of cohesive populations in India, Japan and Mexico.

The analysis with `STRUCTURE` was done for different *K* values (*K* is the number of subpopulations assumed by the model). There is no statistical test available to evaluate the significance of the results for different models, but the most widely used heuristic is the one presented by Evanno[36]. This heuristic shows that the most likely *K* value is *K* = 2, but *K* = 3 and *K* = 6 are also indicated as likely values (Appendix B, Fig. B1). For *K* = 2 and *K* = 3 the partition was the same as with the network approach (Appendix B, Fig. B2). For *K* = 6 the detected partition consisted of five of the six subpopulations detected by the network approach: African, Indian, European, Mexican and East Asian. The Japanese\Chinese division was not detected, but the Masai individuals, assigned to the African subpopulations, were shown to be also likely to belong to a sixth subpopulation (Appendix B, Fig. B2)

### 3.3 SAD analysis

As a demonstration of the SAD analysis, the network with medium threshold (Fig. 1B) was analyzed, and the distribution of the *SA* values for the three communities detected are shown in Figure 2. Figure 3 shows the equivalent analysis of the model-based results using STRUCTURE and assuming three subpopulations (*K* = 3).

**Figure 2:**
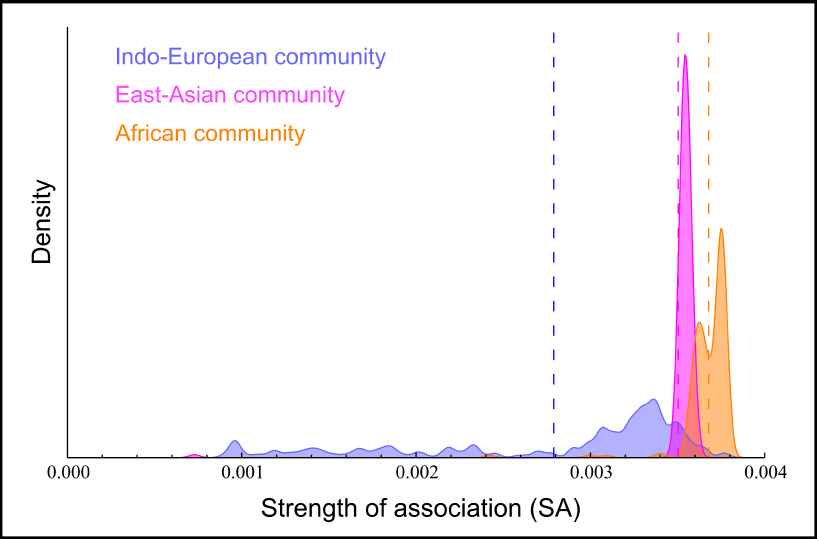
SAD Analysis for the network in Figure 1B. Shown are the distributions of the *SA* values for each of the three communities detected. Mean *SA* for each community is indicated by dashed line.

**Figure 3:**
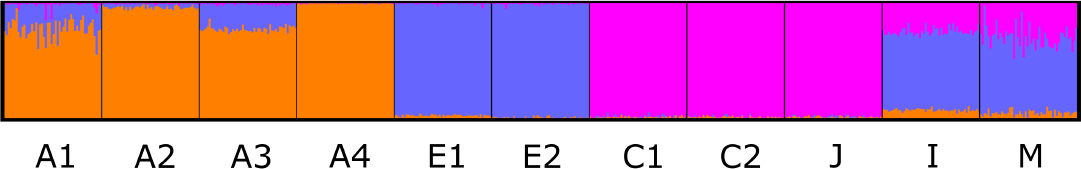
Model-based analysis of human SNP data assuming three subpopulations (*K* = 3)using the program STRUCTURE. The sampled population labels are the same as in Figure 1. The colors of the subpopulations correspond to the colors in Figures 1B and 2.

The SAD of the East Asian community (Fig. 2) has a high mean and is a very narrow distribution, consistent with a subpopulation experiencing limited gene flow. This can be explained from the known historical trend of the relative isolation of East Asia from Europe and Africa.

The SAD of the Indo-European community (Fig.2) is the one with the lowest mean *SA*, and is a wide left-skewed distribution, consistent with a subpopulation with defined core and periphery that experienced extensive gene flow relative to the other subpopulations. Given that the individuals belonging to this community are from European, Indian or Mexican ancestry, a probable explanation is that the core consists of the two European sampled populations and that the Indian and Mexican ancestry individuals have lower association with this group. This can be clearly observed when the distribution is decomposed to three distributions based on the sampled populations (Figure 4).

**Figure 4:**
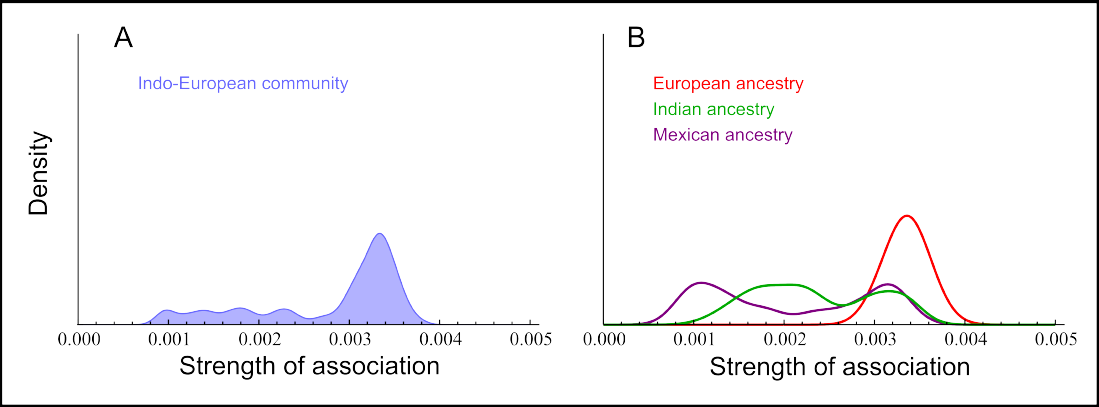
SAD Analysis for the Indo-European (blue) community. (A) Distribution of the Indo-European community as shown in figure 2. (B) *SA* Distribution of the individuals in the community for different sampled subpopulations. It can be seen that the individuals with European ancestry are responsible for the higher *SA* values in the distribution in (A), while the individuals with Mexican or Indian ancestry have lower association to this community.

The distribution of the African community (orange) has a high mean, and is narrow and bimodal. This is consistent with a cohesive subpopulation with limited gene flow, but also that two or more distinct subgroups exist within the population with different levels of association to the community. Figure 5 shows the decomposition of the distribution to the four sampled populations composing it, and it can be seen that the the bi-modality can be explained by the fact that the Masai population (A3) is found to be a distinct population, as detected by some of the community detection algorithms (Appendix A). The `STRUCTURE` analysis also point out to this possibility, as for *K* = 6 it seems that the Masai individuals could possibility be assigned to a different subpopulation, although they are more likely to be assigned to the African subpopulation (Appendix B).

Comparing these results with the results from the model-based analysis, both methods show that the African subpopulation is composed of the same group of individuals (Fig. 1B and Fig. 3). With the model-based analysis, it can be seen that while individuals from Kenya (A2) and Nigeria (A4) have almost no probability to be assigned to other subpopulations, African-ancestry Americans (A1) and Masai individuals (A3) have non-negligible probability to be assigned to other subpopulations, which could be interpreted as that these two groups, while belonging to the African subpopulation, have experienced more gene flow from other subpopulations, mostly from the Indo-European subpopulation. This finding is similar to the one found with the network analysis: these groups are likely to have experienced more gene flow, indicated by the lower mean *SA* of both Masai and African-ancestry Americans than that of the Nigerian and Kenyan groups (Figure 5). However, the *SA* distribution of A1 and A3 are quite different, which implies different evolutionary histories. The A1 SAD is skewed with a long left-tail, indicating that there are a number of African ancestry Americans who are significantly less associated to the community, and are possibly recently admixed individuals. The A3 SAD has a low mean but is symmetric without a wide or skewed tail, possibly indicating that the Masai population has experienced more gene flow, but not in recent times. The recent admixture in the African-ancestry Americans group is consistent with recent higher gene flow experienced by the African ancestry Americans from other American groups.

**Figure 5:**
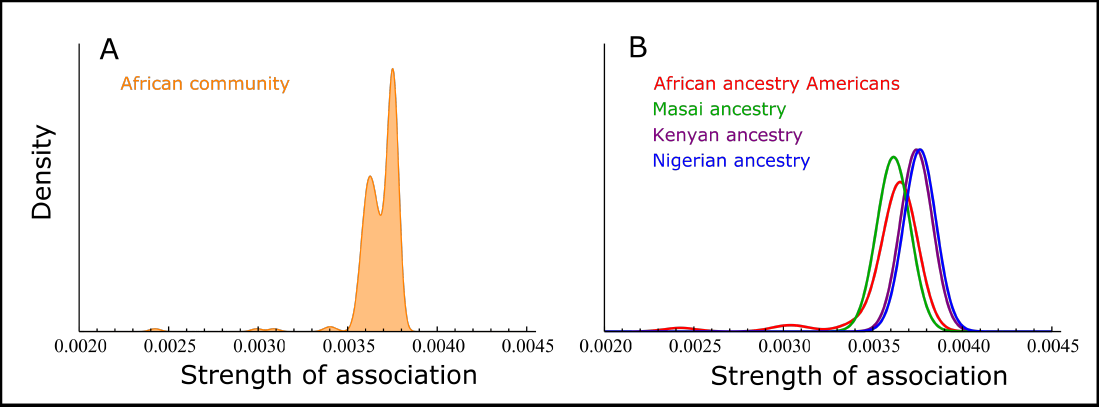
SAD Analysis for the African (orange) community. (A) Distribution of the African community as shown in figure 2. (B) Distribution of the individuals in the community for different sampled populations. The left mode in (A) is due to Masai individuals, which were detected as a distinct population by some algorithms (appendix A). The African ancestry American individuals have slightly lower association to the community than individuals from Nigeria and Kenya, as well as a distinct left-tail, perhaps due to recent admixture with people of European or Native American origin.

## 4 Analysis of simulated data

In order to get a better intuition on the performance of the method, and specifically on the SAD analysis and detection of hybrid and recently admixed individuals, several scenarios were simulated. These simulations consisted of three diploid populations (P1, P2 and P3) of size *N* (2*N* haploids), that have been split *N* generations ago from an ancestral population, also of size *N*. Following the split, migration is assumed to have occurred only between P1 and P2 at rate *m* (proportion of the population per generation). This allowed us to explore the effects of gene flow between P1 and P2 in comparison with an isolated population, P3. Four scenarios with different levels of gene flow between P1 and P2 were examined: no gene flow (S1), low gene flow (S2), medium gene flow (S3) and high gene flow (S4). Table 1 summarizes *m* values for the different scenarios. The coalescence process for 5000 independent SNPs and for *N* = 50 was simulated using the coalescence simulator SIMCOAL2 [37].

### 4.1 SAD analysis of simulated scenarios

For the four simulated scenarios, genetic networks were constructed using equation 3, as done for the human genetic data explained above. Community detection, using the computationally efficient FastGreedy algorithm [38], was performed for edge-removal thresholds starting from 0 in steps of 0.01, until the network was decomposed to many disconnected components. In scenarios S1 and S2, three communities were detected for all thresholds examined, fully corresponding with the three simulated populations (we therefore refer to these communities as P1, P2 and P3). In scenario S3, two communities were detected for low thresholds (0-0.04), fully corresponding to the combined population of P1 and P2 and to population P3 (we refer to these as P1P2 and P3), while for high thresholds (0.05) three communities were detected, fully corresponding to the three simulated populations. For S4, two communities were detected, as for low thresholds in S3. The results are summarized in Table 1, with further details given in Appendix C.

**Table 1:** Strength of Association Distribution (SAD) analysis for simulated scenarios. After the split to three subpopulations, gene flow occurs between P1 and P2 at rate *m*, and P3 remains isolated. Further details are presented in Appendix C.

| Scenario parameters |  |  | SAD characteristics |  |  |
| --- | --- | --- | --- | --- | --- |
| Scenario | Gene flow | $m$ | Detected communities | SAD mean | SAD dispersion |
| S1 | None | 0 | P1, P2, P3 | same for all 3 subpopulations | same for all 3 subpopulations |
| S2 | Low | 0.01 | P1, P2, P3 | P3 has higher mean | same for all 3 subpopulations |
| S3 | Medium | 0.05 | P1, P2, P3<sup>†</sup> | P3 has higher mean<sup>†</sup> | P3 has lower dispersion<sup>†</sup> |
|  |  |  | P1P2, P3<sup>‡</sup> | P3 has slightly higher mean<sup>‡</sup> | same for all 3 subpopulations<sup>‡</sup> |
| S4 | High | 0.1 | P1P2, P3 | P3 has higher mean | same for all 3 subpopulations |

<sup>†</sup>High edge-removal thresholds
<sup>‡</sup>Low edge-removal thresholds

When gene flow was at most medium, the three simulated subpopulations were recovered by community detection, while for high gene flow the procedure was unable to distinguish between P1 and P2. In addition to the detection of subpopulations as the units of structure, the procedure also provided information on hierarchical population structure. For medium gene flow the low edge-removal thresholds P1 and P2 were detected as one community, P1P2, and two separate communities for high edge-removal thresholds. The structure at the two hierarchical levels was detected only at intermediate gene flow levels - with low or high gene flow the same community structure was detected at all thresholds, as structure at one hierarchical level was much stronger and masked the structure at the other level.

The *SA* for all individuals was calculated, and the SAD of the communities was characterized by the mean and Coefficient of Variation (CV), as a measure of dispersion. The analysis was done for networks with high edge-removal thresholds (before network decomposition), but no significant differences were observed for lower thresholds. While for S1 there was no difference in SAD mean of the three detected communities, for the three communities detected in S2 and S3, the mean was significantly higher for P3 (Appendix C, Figure C1). The lower association of individuals from P1 and P2 to their assigned communities, relative to P3, can be attributed to gene flow between these subpopulations. When two communities were detected, the mean for P3 was also higher than the mean of P1P2 (Appendix C, Figure C1), however, in this case, it is not possible to tell if this is a result of gene flow between P1 and P2 or a result of the different number of individuals in P1P2 and P3. Subpopulations represented by a smaller sample size may show higher *SA* values due to the fact that common alleles in these subpopulations may be rare in the entire sample (rare alleles in equation 3 result in edges with higher weights), whereas alleles common in subpopulations with large sample sizes should be relatively common in the entire sample, even if they are rare in other subpopulations.

SAD dispersion may also be an indicator for gene flow, as dispersed *SA* distributions should occur when lineages of many individuals are shared with other subpopulations. While for S2 there was no noticeable difference in dispersion between the communities, for S3 (high threshold) the CV of P3 was significantly smaller than for P1 and P2 (Appendix C, Table C1). For the networks where P1 and P2 were not distinguished (S4 and low thresholds in S3), there was no significant difference in dispersion between the communities. Generally, CV was very low, except for subpopulations experiencing medium gene flow levels, in S2 (Appendix C, Table C1). These results indicate that the high levels of SAD dispersion can be used as an indicator for intermediate levels of gene flow. Overall, the characteristics of the distribution of individual’s *SA*, specifically mean and CV, are correlated with gene flow levels, and allows for the distinction between the four simulated scenarios.

### 4.2 Identifying hybrids and recently admixed individuals

Genetically identifying hybrids or recently admixed individuals of two or more distinct subpopulations is the focus of many studies in ecology and coservation biology [2, 39, 40]. The simulated scenarios were used to test the ability of SAD analysis to identify hybrids (offspring of individuals from two different subpopulations) and recent descendants of hybrids. In each scenario, a first-generation hybrid, H1, was simulated by randomly selecting one individual from P1 and one individual from P2, and generating a random offspring from the parents’ genotypes. Similarly, H1 was backcrossed with a randomly selected individual from P1 to produce a second-generation hybrid, H2. These individuals were then added to the simulated population, and the SAD was analyzed to see whether the hybrid individuals can be identified as outliers (the analysis was done separately for H1 and H2). This process was repeated 20 times in all scenarios where the three subpopulations were identified (S1, S2, and high threshold of S3).

As all repeated simulations showed similar results qualitatively, we show only one example of the outlier identification process (Figure 6). For S1 and S2, both H1 and H2 were easily identified as outliers with noticeably lower SA values than the rest of the individuals in their assigned community. For S3, H1 was revealed as a hybrid with lower SA values, but H2 was not.

**Figure 6:**
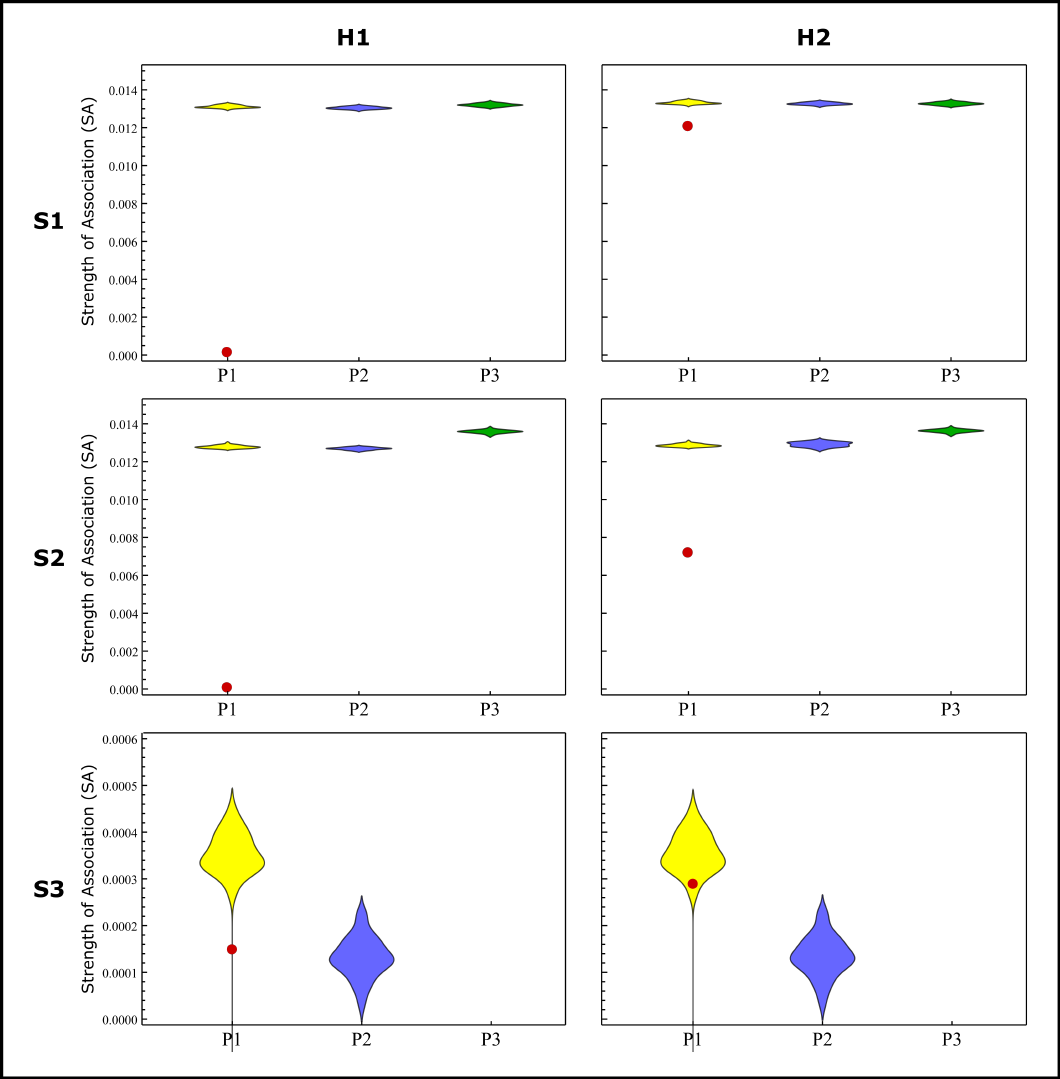
An example of hybrid identification using SAD analysis. Distribution charts show the SADs for the three detected communities, with the *SA* of the hybrid shown as red dot (distribution of P3 in S3 is out of scale). The analysis was performed on simulated scenarios S1, S2 and S3, where H1 is a hybrid between populations P1 and P2, and H2 is a second-generation hybrid between H1 and P1. Hybrids are identified as low SAD outliers in all cases except for H2 in S3.

In all cases when H1 or H2 were identified, equation 2 was examined to determine which k minimizes the equation. This was done in order to identify the second subpopulation that the hybrid individual is associated with. The second population of origin was identified for H1 in all scenarios, while for H2 it was recovered only for low levels of gene flow. Further details regarding identification of hybrid individuals and more detailed results appear in Appendix C. The results of these simulations show that outliers of *SA* distributions may be considered as potential hybrids, but the ability of the method to identify such individuals decreases with decreasing differentiation between the populations in question, as well as with the number of generations after the hybridization event.

## 5 Discussion

We present a model-free distance-based approach for analysis of population structure at the individual-subpopulation level, which does not entail the assumptions of an underlying model or any prior conditions. The approach is set in a network theory framework and uses the concepts of *community* and *modularity*, which allows for computationally efficient assignment of individuals to sub populations. The computational efficiency makes this approach applicable also in cases where many loci are studied. An additional exploratory analysis of the *SA* distributions of the communities can be used to study population structure beyond assignment of individuals to populations, by evaluating the strength to which individuals are associated with their assigned subpopulations. This may be useful to detect hybrid and recently admixed individuals, as well as to explore finer details of the population structure, as demonstrated by analysis of simulated and real data sets.

Clustering individuals based on genetic composition has become important for many studies in various fields such as ecology, conservation, medicine and anthropology, and there are now several methods and programs addressing this task[9, 41]. The more developed approach is the model-based approach, which utilized Bayesian or maximum-likelihood methods to estimate allele frequencies of *K* hypothetical populations, under model assumptions such as HWE, and assign probabilities of assignment for individuals to these populations. The alternative, model-free, approach is based on analysis of distance (dissimilarity) or similarity matrices, which summarize some measure of distance or similarity between all pairs of individuals. These matrices are usually analyzed using multidimensional scaling (MDS) such as PCA, and projected to two or three coordinates for visualization (e.g. `PLINK`[42], `EIGENSOFT`[43], `GENEALEX`[44, 45]). Clusters are determined either visually or using other clustering methods, such as k-means[46], often requiring a priori definition of the number of clusters to be found, *K*. Similarly, other distance-based methods apply spectral clustering techniques and analyze the eigenvalue spectra of similarity matrices ([47]). Since model-based and distance-based approaches differ significantly in methodology and prior assumptions, researchers often apply both approaches to the same data set in order to ensure that interpretations are not biased by methodology.

The network approach we present here falls under the distance-based clustering category, and no underlying model is assumed. Clustering is done by locating groups of nodes that are strongly connected within the group but weakly connected to nodes outside the group. The strength of the intra-community connections is not defined by a fixed value, but rather is determined relative to the network structure. For example, spectral community detection methods [31, 48] do not analyze the spectra of the similarity matrix itself, but rather address the modularity matrix which is constructed by comparing the strength of connection between nodes with what would be expected in a random network with similar structural properties (i.e. same node degrees). Community detection methods, used in our approach, therefore differ from other distance-based clustering methods in what is considered a “good cluster”, and have been shown to detect meaningful clusters in many systems where clusters are better defined by relative relations between elements than by other distance-based definitions[26, 27]. In this sense, community detection is more similar to model-based approaches, but here no prior assumptions are required. This also makes it more natural to weigh pairwise genetic similarity by allele frequencies to strengthen similarity between individuals sharing rare alleles relative to those sharing common alleles (equation 3), whereas other distance-based methods more commonly employ only identity-by-state (IBS) measures to define similarity or distance between individuals.

Ideally, population-level studies would benefit from exploring structure using our network approach in combination with model-based clustering methods and MDS visualization, as these complement each other and may give a more robust and detailed picture. In the example analyzed in this paper using human SNP data, the model-based analysis and the network analysis were relatively in agreement regarding assignment of individuals to subpopulations. However, the network analysis did detect the difference between the Chinese and Japanese groups which was not detected by the software `STRUCTURE` Additionally, the SAD analysis revealed differences in gene flow experienced by the Masai and African-ancestry American groups, but these groups appear very similar in the `STRUCTURE` analysis (Figure 3).

The `STRUCTURE` analysis did indicated biologically-meaningful structure at different hierarchical levels, however this was done using a heuristic, and the hypothesis of population structure at any hierarchical level is not statistically testable. Statistically testable hierarchical partitioning of population structure is, of course, feasible and common in subpopulation-population level methods (e.g. F-statistics, AMOVA), but is not common in individual-subpopulation procedures, where putative subpopulations are not defined a priori. With the approach presented here, a similar hierarchical structure was detected through systematic examination of edge-removal thresholds, but here each hierarchical level was statistically tested and shown to be significant. Statistically testing hierarchical structure is important since, otherwise, analyses are forced either to recover only the strongest level, missing biologically-relevant information about the system, or report many hierarchical levels, without a procedure for determining which are meaningful and which are not. With the network method presented here, different significant community structures emerge, producing a semi-hierarchical structure, in the sense that a community partition at a given level does not depend on “higher” level partitions. True hierarchical community partition procedures[49, 50] can possibly be useful for hierarchical population structure analysis, but in most of these procedures each level is constrained by higher levels.

Another related issue, that has been a concern in model-based implementation, is the assessment of the number of subpopulations, *K*[7, 36], as *K* is usually one of the model parameters. This problem arises also when applying clustering techniques on results of distance-based methods, such as k-means on MDS results. By setting *K*, these procedures regard the subpopulations as equivalent, even though this is often not the case. For example, for the network shown in Figure 1B, *K* = 3, however the three subpopulations show very different distributions of within-population relatedness (Fig. 2). In the network-based approach there is no such issue, as most community detecting methods do not a priori assume *K*, but rather find the optimal *K* that maximizes modularity (e.g. [32, 38]), or acquires *K* as part of the detection process (e.g. [31, 51, 52]), without assuming any equivalence of the communities. This is due to the fact that modularity is a trusted and well-studied measure for the quality of community partitions[32].

With whole genome sequencing becoming more and more accessible, procedures for population structure analysis must also take into account computational considerations. The procedure presented here is composed of three consecutive steps, with construction of the network taking *O*(*Ln*^2^) time, where *n* is the number of individuals in the sample and *L* is the number of loci involved in the analysis. Computation time for community detection depends on the algorithm used, but fast near-linear algorithms, taking *O*(*n + m*) time, where *m* is the number of edges in the network, and approaching *O*(*n*) time for sparse networks (which can be constructed using high enough edge-removal thresholds), are already available[51]. SAD analysis depends on *m* and on the number of communities detected, *c*, and takes *O*(*cnm*) time, approaching *O*(*cn*^2^) for sparse networks. Only the first step involves *L*, therefore the computation time of the entire procedure is linear with respect to the number of loci, and there should be no computational limitation for including full genome sequences in analyses.

Since the genetic-similarity measure, the threshold and the community detection algorithm remain, for now, used-defined, and may result in different population structures, care must be taken when defining these parameters, and preferably several options should be explored. Further studies may provide guidelines for setting these parameters as a function of the particulars of the system under study. Network theory, and particularly community detection, is a highly active field of research, but our understanding of the usefulness of particular community detection procedures to different types of networks is still minimal, and future advancements in network theory may provide clearer guidelines for algorithm and threshold choice. As with most other individual-subpopulation level clustering methods, significant linkage disequilibrium (LD) between loci should be avoided, and data should be screened to ensure that loci can be regarded as independent. The extent to which LD will affect both community partition and the significance test presented here should also be explored in the future.

We believe a network approach may provide an additional complementary viewpoint on population structure analysis, one less hampered by prior assumptions. Moreover, defining population genetic problems in network terminology is important for establishing the usage of network theory in population genetics. Currently, many tools and methods are developed within network theory in order to study complex systems, and these methods may become accessible to the field of population genetics once network terminology is appropriately incorporated in population genetic theory and practice.

The method presented here is implemented in the program NetStruct, which uses community detection algorithms implemented in the software package igraph[35], and is available at https://github.com/GiliG/NetStruct.

## Acknowledgments

The authors would like to thank Yair Zarmi, Sharon Renan, Naama Shahar and Shai Pilosof for insightful comments and discussions. This research was supported by United States-Israel Binational Science Foundation grant 2011384 awarded to SB, ART and Amos Bouskila. This is publication XXX of the Mitrani Department of Desert Ecology.

